# Sparse Binary Relation Representations for Genome Graph Annotation

**DOI:** 10.1101/468512

**Authors:** Mikhail Karasikov, Harun Mustafa, Amir Joudaki, Sara Javadzadeh-No, Gunnar Rätsch, André Kahles

**Affiliations:** Department of Computer Science, ETH Zurich, Zurich 8092, Switzerland; University Hospital Zurich, Biomedical Informatics Research, Zurich 8091, Switzerland; SIB Swiss Institute of Bioinformatics, Lausanne 1015, Switzerland

**Author notes:** http://bmi.inf.ethz.ch.

**Keywords:** sparse binary matrices, binary relations, genome graph annotation, compression

## Abstract

High-throughput DNA sequencing data is accumulating in public repositories, and efficient approaches for storing and indexing such data are in high demand. In recent research, several graph data structures have been proposed to represent large sets of sequencing data and to allow for efficient querying of sequences. In particular, the concept of labeled de Bruijn graphs has been explored by several groups. While there has been good progress towards representing the sequence graph in small space, methods for storing a set of labels on top of such graphs are still not sufficiently explored. It is also currently not clear how characteristics of the input data, such as the sparsity and correlations of labels, can help to inform the choice of method to compress the graph labeling. In this work, we present a new compression approach, *Multi-BRWT*, which is adaptive to different kinds of input data. We show an up to 29% improvement in compression performance over the basic BRWT method, and up to a 68% improvement over the current state-of-the-art for de Bruijn graph label compression. To put our results into perspective, we present a systematic analysis of five different state-of-the-art annotation compression schemes, evaluate key metrics on both artificial and real-world data and discuss how different data characteristics influence the compression performance. We show that the improvements of our new method can be robustly reproduced for different representative real-world datasets.

## 1 Introduction

Over the past decade, there has been an exponential growth in the global capacity for generating DNA sequencing data [22]. Various sequencing efforts have started to amass data from populations of humans [1] and other organisms [23,24]. For these well studied organisms, already assembled reference sequences are the common starting point for comparative and functional analyses. Unfortunately, a large proportion of DNA sequencing data, in particular data originating from non-model organisms or collected in metagenomics studies, are lacking a genome reference. Whereas general guidelines exist for the former case [8], genome assembly for metagenomics is much less well defined. Its vastness and the currently lacking standards for indexing such data make an integrated analysis daunting even for field experts.

To make this host of data efficiently searchable, it is necessary to employ a search index. However, when building an index on the sequence data alone, only presence or absence of a query can be tested. To support relating queries to information such as source genomes, haplotypes, or functional annotations, additional labels must be associated with the index. To facilitate this, approaches for storing additional data on an indexed graph have been suggested, such as the gPBWT [16] for storing haplotype information as genome graphs or succinct representations of *labeled* de Bruijn graphs [13,11,4] for the representation of sets of sequences. In this context, dynamic representations of such data have also recently received attention [14,19].

The problem of efficiently representing these types of relations is also addressed in other fields. Commonly referred to as *compressed binary relations*, a growing body of theoretical work addresses such approaches [6]. Successful applications of similar techniques include the efficient representation of large web-graphs [7] and RDF data sets [5]. We will provide a more detailed description of some of these approaches in Section 2.

In this work, we present a new method for compressing abstract binary relations. Providing as background a comprehensive benchmark of existing compression schemes, we show that our approach has superior performance on both artificial and real-world data sets.

Our paper has the following structure: After introducing our notation, we begin by defining the abstract graph, and the associated annotation structures that we wish to compress (Section 2.1). We then provide descriptions of our proposed compression technique and competing methods (Section 2.2). Finally, we compare the compression performance of these methods on different types of graph annotations (Section 3) and close with a brief discussion of our results and an outlook on future work (Section 4).

## 2 Methods

After introducing our notation, we will give an overview of all methods implemented for this work and provide a description of our methodological contributions.

### 2.1 Preliminaries

We will operate in the following setting: We are given a *k*-dimensional de Bruijn graph over a set of given input sequences *S*. The node set *V* shall be defined as the set of all consecutive sub-sequences of length *k* (*k*-mers) of sequences in *S*

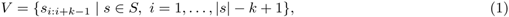

where *s*_*i*:*j*_ denotes the sub-sequence of *s* from position *i* up to and including position *j*, and *|s|* is the length of *s*. A directed edge exists from node *u* to node *v*, if *u*_2:*k*_ = *v*_1:*k*−1_.

In order to represent relations between sources of the input sequences *S* and the nodes *V*, we now define the concept of a *labeled de Bruijn graph* and proceed by discussing the more general problem of representing a graph labeling.

Each node *v* ∈ *V* that we refer to as an *object* is assigned a finite set of labels *ℓ*(*v*) *⊂ L*. We represent this *graph labeling* as a binary relation 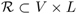. A trivial representation of 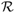 taking *|V | ⋅ |L|* bits of space is a binary matrix *A* ∈ {0, 1}^|*V* |×|*L*|^. We will use *A*^*i*^ and *A*_*j*_ to denote its rows and columns, respectively.

In the following sections, we will discuss various methods described in the recent literature and present our improvements in efficiently representing 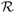. In addition to minimal space, we also require that the following set of operations can be carried out efficiently on the compressed representation of 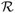:

`query labels`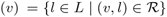 Given an object *v* ∈ *V* (a *k*-mer in the underlying de Bruijn graph), return the set of labels *ℓ*(*v*) assigned to it.

`query objects`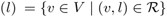 Given a label *l* ∈ *L* (e.g., a genome or sample ID), return the set of objects assigned to that label.

`query relation`(*v, l*) Given an object *v ∈ V* and a label *l ∈ L*, check whether (*v, l*) is in the relation 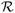, `query relation`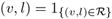.

### 2.2 Binary Relation Representation Schemes

For compressing the binary relation 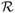, we consider the following representations suggested in recent literature. As an abstraction, we will use the representation of 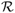 as a binary matrix *A* ∈{0, 1} ^*|V*|×|*L*|^ (referred to as the *binary relation matrix*) to illustrate the individual methods.

#### Column-major Sparse Matrix Representation

As a simple baseline technique, we compress the positions of the non-zero indices in each column independently using Elias-Fano encoding [17]. While this method does not take into account correlations between columns for compression, this feature allows for a trivial parallel construction implementation in which each column is computed in a separate process. For our experiments, this serves as the initial representation of the binary matrix, which is then queried during the construction of all other matrix representations.

#### Flat Row-major Representation

As a second baseline method, this representation concatenates all rows of *A* into a joint vector that is subsequently compressed using Elias-Fano encoding. This approach, for instance, is used by VARI [13] and its extensions [3].

#### Rainbowfish

The current state-of-the-art for genome graph labeling is a row-major representation of the binary relation matrix *A* in which an optimal coding is constructed for the set of rows in *A* [4]. More precisely, let 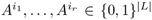 denote the unique rows of *A*, sorted by their number of occurrences in *A* in non-increasing order, where *r* ≤ |*V* |. To encode *A*, we start by forming a matrix *A*′ ∈ {0, 1}^*r*×|*L*|^ of sorted unique rows, 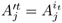. Then we compress *A′* with the flat row-major representation using an RRR vector (named after the initials of the three original authors [20]) as the underlying storage technique and construct a *coding vector* (*i*(*v*) − 1)_*v*∈*V*_, where *i*(*v*) maps each node *v* ∈ *V* to the index of the row in *A′* corresponding to the labeling of *v*. The coding vector is represented in a variable-length packed binary coding with a delimiter vector [4] compressed into an RRR vector [20].

#### Binary Relation Compressed with Wavelet Trees (BinRel-WT)

This method involves a translation of the 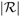 non-zero elements of *A* into a string, which is then represented using a conventional wavelet tree [6]. Given the binary relation matrix, its set bits are iterated in row-major order and their respective column indices are stored contiguously in a string over the alphabet{1*,…*,|*L*|} represented with a wavelet tree that enables efficient queries. The numbers of set bits in each row of *A* are stored in a delimiter vector using unary coding and compressed into an RRR vector.

#### Hierarchical Compressed Column-major Representation (BRWT)

Described as Binary Relation Wavelet Trees (BRWT) in the original literature [6], in contrast to BinRel-WT, this representation directly acts on binary matrices without translation into a sequence. First, an index vector *I* with elements 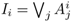 is computed by merging all matrix columns through bitwise-OR operations on the rows and stored to represent the root of the tree. Then, the rows composed entirely of 0s are discarded from *A* and two equal-sized submatrices *A′* and *A′′* (which may contain rows composed entirely of 0s) of the binary relation matrix *A* are constructed by splitting *A* and are passed to the left and right children of the root. The compression proceeds recursively. Construction terminates when a node is assigned a single column, which is stored as its index column (see Figure 1a)). For reconstruction of the matrix elements, it is sufficient to only store the index vectors associated with each node of the BRWT tree.

**Fig. 1.**
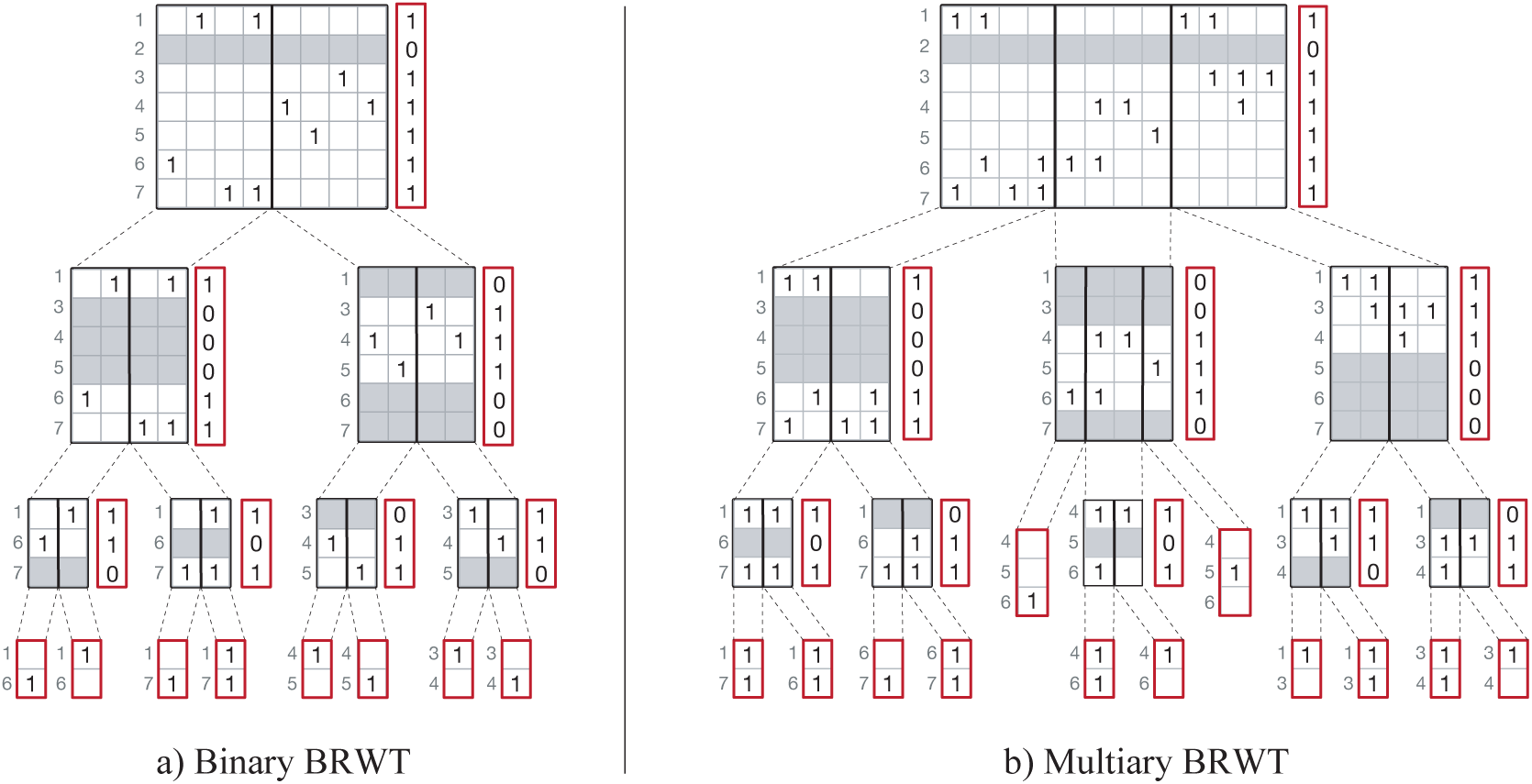
Schematic of hierarchical compressed column-major representations — a) BRWT for the binary case. Grey rows correspond to all-zero rows, also indicated through the vector to the right of each matrix. Each child encodes only non-zero rows of the submatrix passed to it by its respective parent. Numbers to the left of each matrix are the respective row-indices in the initial matrix. b) Multi-BRWT in the multiary case. Notation is as in the binary case. Stored vectors are shown in red.

In the next section we consider the problem of topology optimization during BRWT tree construction and propose *Multi-BRWT*, an extension of BRWT that allows its nodes to have arbitrary numbers of children as well as to arbitrarily distribute the columns of a parent node to its children. Afterwards, we propose a two-step approach for Multi-BRWT construction along with two specific algorithms as its implementation for improving the compression performance of Multi-BRWT.

### 2.3 Multiary, Topology-Optimized BRWTs

Our first extension to the BRWT scheme is the introduction of an *n*-ary tree topology, Multi-BRWT (Split *n*), allowing for matrices to be vertically split into more than two submatrices (see Figure 1b)). The construction and querying for Multi-BRWT (Split *n*) is analogous to the case of binary BRWT. In computational experiments on artificial and real-world data we show that in most cases, Multi-BRWT (Split *n*) with arity greater than two provides a higher compression ratio than the simple binary BRWT scheme (see Section 3). Note that Multi-BRWT (Split *n*) with the maximum allowed arity *n* = |*L*| is equivalent to the baseline column-major sparse matrix representation as it keeps all columns of the input binary relation matrix unchanged except for the case when the input matrix has all-zero rows.

To proceed with our second extension, let us consider binary relations with far fewer labels than objects, |*L*| ≪ |*V*|, a condition that is commonly met in annotated genome graphs from biological data. In these contexts, the number of *k*-mers is usually in the billions and the number of labels is on the order of thousands (see Section 3.2).

Our second extension consists in introducing arbitrary assignments of columns from the matrices encoded in the nodes of the Multi-BRWT to their children. These assignments are represented by dictionaries stored in the Multi-BRWT nodes, but the |*L*| ≪ |*V*| constraint makes the space overhead from storing these negligible compared to the space needed to encode the index vectors. Thus, we exclude the problem of representing these assignments from further consideration and leave that as a small technical detail.

We now focus on the problem of constructing a Multi-BRWT tree structure that satisfies certain local optimality conditions with respect to compression ratio.

#### Problem Setting

Let us set this problem formally. Given a binary matrix *A*, let *𝒯* be the set of all Multi-BRWT trees representing *A* (i.e., the set of all rooted trees with |*L*| labeled leaves). Let *Size*(*I*) denote the size of a compressed binary vector *I* in bits. For instance, if *I* is of length *n* with *m* set bits, *Size*(*I*) = *n* for an uncompressed bit vector, and 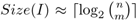 for RRR vectors [20]. We then neglect the space required for dictionaries defining the column assignments and we define the size of the Multi-BRWT tree *T* ∈ *𝒯* as the space required to store all its index vectors including the vectors in leaves:

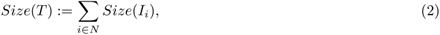

where *N* is the set of all nodes of the Multi-BRWT tree *T* and *I*_*i*_ corresponds to the index vector stored in node *i*. Thus, we wish to find an optimal Multi-BRWT tree by minimizing the storage space,

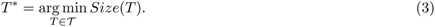

We will refer to this as the Multi-BRWT problem.

#### Optimized Multi-BRWT Construction

By analogy to the NoSQL table compaction problem [9], it can be shown that Multi-BRWT constrained on the space of binary trees with the uncompressed bit vector representation as the underlying structure for storing the index vectors is NP-hard. Thus, we propose a two-step approach for finding a good Multi-BRWT structure (see Figure 2). First, we build a binary Multi-BRWT tree by hierarchical clustering of the index vectors according to their similarity, the number of shared set bits. Then, we optimize the arity of the chosen Multi-BRWT by selecting a node subset *N′* which includes the root and leaves of the base Multi-BRWT tree, {*r*, *v*_1_, …, *v*_|*L*|_} ⊂ *N′* ⊂ *N*. To keep the resulting Multi-BRWT tree valid (allowing for reconstruction of the initial matrix *A*), we reassign all nodes in *N′* to their nearest common ancestors remaining in *N′*.

**Fig. 2.**
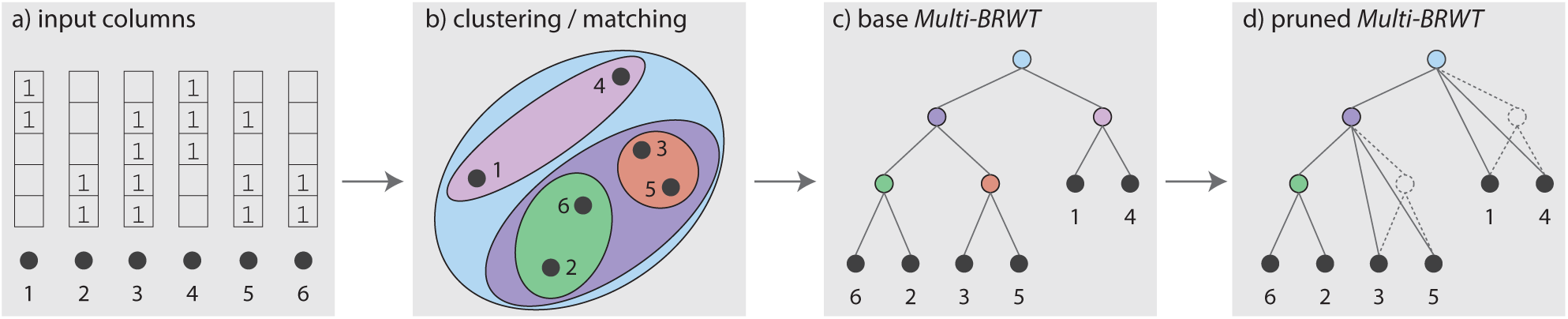
Schematic describing the construction of *Multi-BRWT* — a) The columns of the input binary matrix depicted as numbered black dots are considered independently. b/c) Columns are hierarchically pair-matched based on number of shared entries, forming the base Multi-BRWT topology. d) Pruning internal nodes of Multi-BRWT to optimize the tree structure for a smaller representation size.

As a specific implementation of the proposed two-step construction approach, we consider two heuristic greedy optimization procedures. In the first step, we perform greedy matching of the index vectors starting from the columns of the input binary relation matrix, and repeat recursively for the aggregated parent index vectors until we merge all into a single index vector placed in the root. In the second step of the construction approach, we consider another greedy algorithm for optimizing the size of the Multi-BRWT tree by removing some of its internal nodes and thereby increasing the arity of the tree.

##### Greedy Pairwise Matching for Finding a Base Multi-BRWT Approximation

To find an initial approximate solution to the Multi-BRWT problem (base Multi-BRWT), we propose a greedy algorithm in which an initial greedy pairwise matching (GPM) step is performed on the columns of the input binary relation matrix *A* to optimize their initial order prior to construction (see Figure 2a–c)). Given the input columns *A*_1_, …, *A*_|*L*|_ and their corresponding object queries *o*_*i*_ = `query objects`(*i*), we first compute cardinalities of their pairwise intersections *s*_*ij*_ = |*o*_*i*_ ∩ *o*_*j*_|. Then, we sort all the computed similarities {*s_ij_*} in non-increasing order and match pairs of columns greedily. Afterwards, we compute the aggregated index columns by merging the matched columns through bitwise-OR operations to form the index vectors and repeat this algorithm recursively.

##### Efficient Pairwise Distance Estimation

The proposed greedy approximation method takes as input a matrix of pairwise column similarities {*s*_*ij*_}. For *m* input columns of length *n*, computing each entry of this matrix costs 𝒪(*n*), and thus, the time complexity of computing the full similarity matrix is 𝒪(*nm*^2^), which is a considerable overhead for datasets with a typical size of *m* ~ 10^3^ and *n* ~ 10^9^. To make the estimation of the pairwise similarities cheaper, we approximate these on a submatrix composed of rows sampled randomly from matrix *A*. Moreover, we prove the following lemma to show that using just 𝒪(log(*m*)/*ε*^2^ random rows is sufficient for approximating the pairwise similarities with a small relative error *ε* with high probability, if each column has a sufficiently large number of set bits.

#### Lemma 1 (Subsampling lemma)

*Suppose we are given subsets of a universe set, o*_1_, …, *o*_*m*_ ⊂ {1, …, *n*}, *with the minimum cardinality 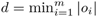, *d* > 0. We sample the elements of {*1, …, *n*} *independently with the same probability p, to form a sampled set of objects S ⊂ {*1*, …, n} and define subsampled sets as 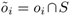*. Consider the union cardinalities u_*ij*_ = *|o*_*i*_ ∪ *o*_*j*_| *with their approximators 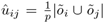. For all 0 < ε < 1, 0 < δ < 1, and*

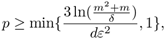

we claim

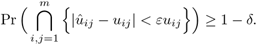

See Supplementary Section 2 for proof.

According to Lemma 1, with the subsampling technique we can approximate the union cardinalities up to an *ε*-fraction with high probability. Similar bounds can be obtained for sufficiently large intersection cardinalities, estimated in the proposed greedy pairwise matching algorithm.

##### Refining Multi-BRWT by Pruning

Starting the procedure in the leaves’ parents and applying it to each node except for the root recursively, we estimate the cost of removing each current node by the following formula

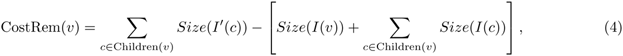

where *I*(*v*) denotes the index vector stored in the node *v* and *I′*(*c*) denotes the updated index vector that would be stored in the node *c* if its parent *v* was removed and the node *c* was reassigned to its grandparent. Now we simplify the formula for estimating the cost of removing a node in Multi-BRWT by introducing an assumption that the size of bit vector *I* of length *n* with *m* set bits is fully defined by these two parameters, i.e. *Size*(*I*) = *Size*(*n, m*). Now, it is easy to see that after reassigning the node *c* with the index vector *I*(*c*) of length *n*_*c*_ with *m*_*c*_ set bits to the parent of its parent *v* with index vector *I*(*v*) of length *n*_*v*_ with *m_v_* set bits, the node *c* updates and replaces its index vector *I*(*c*) with a vector *I′*(*c*) of length *n*_*v*_ with *m*_*c*_ set bits. This provides us with the following simplified formula for estimating the cost of removing a node from the Multi-BRWT tree

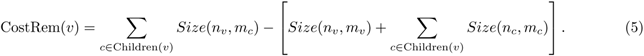

Formula (5) can be efficiently computed without rebuilding the current structure of the Multi-BRWT. As a result, a decision about removing node *v* from the Multi-BRWT is made if the cost CostRem(*v*) is negative, leading thereby to a decrease of the Multi-BRWT in size. In our practical implementation we use the following formula for approximating the size required for storing an RRR bit vector [15] with block size 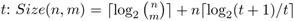.

### 2.4 Implementation Details

We implement the underlying de Bruijn graph as a hash table storing *k*-mers packed into 64bit integers with 64bit indexes assigned to the *k*-mers, or as a complete de Bruijn graph represented by a mapping of *k*-mers to 4^*k*^ row indexes of the binary relation matrix.

In the column-major representation, the columns of the binary relation matrix are stored using SD vectors implemented in `sdsl-lite` [10]. The same data structure is used for storing the single long vector in the row flat representation.

BinRel-WT (sdsl) compressor uses the implementation of wavelet tree from the `sdsl-lite` library, using an RRR vector to store its underlying bit vector. The delimiter vector uses the RRR vector implementation from `sdsl-lite`.

The BinRel-WT compressor uses the binary relation implementation from https://github.com/dieram3/binrel_wt. This implementation stores the underlying bit vector of the wavelet tree in uncompressed form.

Our BRWT is implemented as a tree in memory, compressing the index vectors as RRR vectors. To avoid multiple passes through the matrix rows, we construct the BRWT using a bottom-up approach. Given a fixed clustering of the matrix columns, the leaves of the BRWT are constructed first, followed by their parents constructed for the index vectors propagated from the children nodes. To speed up the greedy matching algorithm, we sample randomly 10^6^ rows in each experiment and use those to approximate the number of bits shared in the input columns and the index vectors during the Multi-BRWT construction. When optimizing the tree arity (as described in Section 2.3), we use the formula 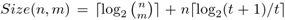 as an estimate for the size of bit vector *I* of length *n* with *m* set bits, which is provided by the authors of *sdsl-lite* for the implementation of RRR vectors [10]. We use a block size of *t* = 63.

All SD vectors are constructed with default template parameters, while all RRR vectors are constructed with a block size of 63.

#### Code Availability

All methods implemented and evaluated in this paper are available at https://github.com/ratschlab/genome_graph_annotation.

### 2.5 Data

#### Simulated Data

To profile our compressors, we generated several different series of synthetic binary matrices of varying densities (see Supplementary Section 1 for a more detailed description). In total we generated three different kinds of series: i) random matrices with uniformly distributed set bits, ii) initially generated random matrix rows duplicated and permuted randomly, iii) initially generated random matrix columns duplicated and permuted randomly. The motivation behind these series is as follows: The best performing state-of-the-art compressors exploit redundancy between rows of the binary relation matrix [19]. However, the usual structure of annotated de Bruijn graphs often implies a correlation structure on the columns not necessarily leading to redundant rows, for instance when the sequences of many similar or closely related samples are inserted. While for a small (and sufficiently highly correlated) number of columns this correlation translates into rows and increases the number of redundant ones, for larger label sets this is usually not the case. Thus, approaches exploiting correlation structure on the columns might fare better. To test this hypothesis, we generated three different kinds of synthetic data, reflecting uncorrelated rows/columns, redundant rows, and redundant columns for series i), ii), and iii), respectively. Please note, that approach ii) is the most favorable for the state-of-the-art, as row redundancy rather then high correlation is simulated.

#### Real-World Data

For evaluating all approaches in a real-world setting, we have chosen two data sets well-known in the community and representative of typical applications.

##### Kingsford Human RNA-Seq

This dataset consists of 2,652 Human RNA-Seq experiments originally drawn from [21] and subsequently used in [19] for comparison.

##### NCBI RefSeq

This dataset consists of all 79,448 reference sequences from Release 88 of the NCBI RefSeq database [18]. Each sequence has been annotated with its associated `family` rank taxonomic ID from the NCBI Taxonomy [2]. This results in a total of 3,173 unique labels for the sequences.

## 3 Results and Discussion

### 3.1 Experiments on Artificial Data

Based on the artificial dataset described in Section 2.5, we evaluated how the compression performance changes depending on the characteristics of the input binary relation matrix *A* of a simple structure.

#### Dependency of Compression Ratio on Matrix Structure

One of the key characteristics of the binary relation matrix *A* is its density, the number of set bits divided by the total number of entries in *A*. For reference, the labels for a sequencing-based de Bruijn graphs typically exhibit very low densities, commonly *<* 0.5%. Especially in this low-density region, we find that the properties of the binary relation matrix have a strong effect on the compression ratio of individual methods. A second determinant of performance is whether any assumptions are made on the properties of the data.

On sparse, fully random data, the baseline compressors fare very well (Figure 3a)), as no assumptions can be made about relationships. Notably, Rainbowfish, which exploits redundancy among the rows, generates considerable overhead for very low densities. In the field of BRWT methods, the Multi-BRWT is closest to the best performing choices.

**Fig. 3.**
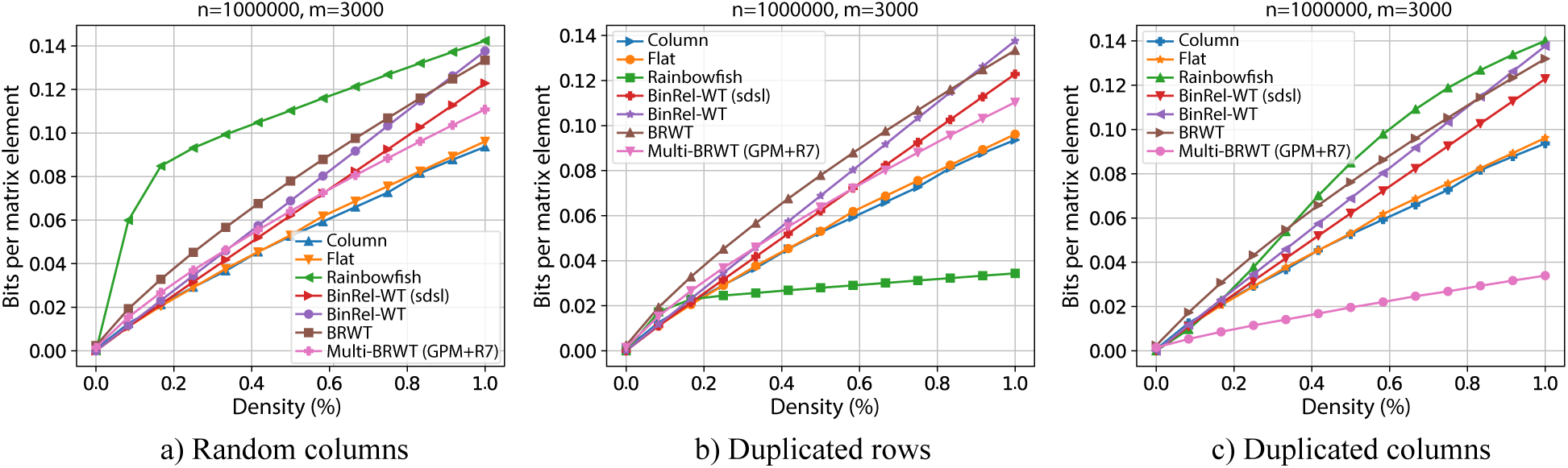
Size of the representation of *A* ∈ {0, 1}^10^6^×3⋅10^3^^ with densities *d <* 0.01 using different approaches: a) uniformly random bits; b) uniformly random rows with multiplicity 5; c) uniformly random columns with multiplicity 5. We expect approach c) to be best reflecting the real-world data of a de Bruijn graph built on related sequences.

In the setting of redundant rows (data set ii); Figure 3b)), as expected, Rainbowfish shows the strongest performance, clearly exploiting the row redundancy. Again, among the BRWT methods the Multi-BRWT performs best.

Finally, in the setting that comes closest to a typical task of labeling de Bruijn graphs derived from sequencing data (Figure 3c)), the Multi-BRWT approach shows superior performance. Exploiting the correlated columns of the matrix, Multi-BRWT achieves a 5-fold improvement in compression ratio compared to Rainbowfish and more than 2-fold compared to the closest competitor. Notably, the baseline binary BRWT has no advantage over the other baseline methods. Further, we observe that this performance gain increases with the total number of columns in the matrix (Supplemental Figures 1 and 2).

### 3.2 Experiments on Real-World Data

To compare the compression performance of the considered methods under a variety of conditions, we have constructed two test datasets that exhibit different matrix sparsity characteristics.

#### Kingsford Human RNAseq (2,652 read sets)

We filtered the 2,652 raw sequencing read sets with the KMC [12] tool to extract frequent unique *canonical k*-mers (defined as the lexicographical minimum of the *k*-mer and its reverse complement) from each (*k* = 20). We used the same *k* value and thresholds for the *k*-mer frequency level as [19]. Using the *k*-mers extracted, we constructed a de Bruijn graph with 3,693,178,415 nodes and annotated these with their source read sets, which resulted in 2,586 labels (66 filtered read sets were empty) and a binary relation (annotation) matrix of density ~ 0.19%. As a baseline for comparison, we used the straightforward column-compressed annotation, which required a total of 36.56 Gigabytes (Gb) of space. We used this as a starting point to convert the annotation into the other formats.

The results are summarized in Table 1. As expected, the simple row-based and BinRel-WT representations require more than 30Gb in total. The current state-of-the-art method, Rainbowfish, reduces this by 23% to 23.16Gb, exploiting the redundancy of rows in the input matrix. The basic BRWT benefits from the column correlation and drastically improves on Rainbowfish, showing a 39% lower size. We further reduce this size through our generalized approach using Multi-BRWT. While some increase in arity reduces size compared to the binary case, a higher arity does not necessarily translate into lower space, as certain submatrices do not benefit from being grouped. The smallest fixed-arity representation is Multi-BRWT (Split 5), requiring 13Gb of storage space and 222 minutes of compute time to construct with four threads.

**Table 1.**
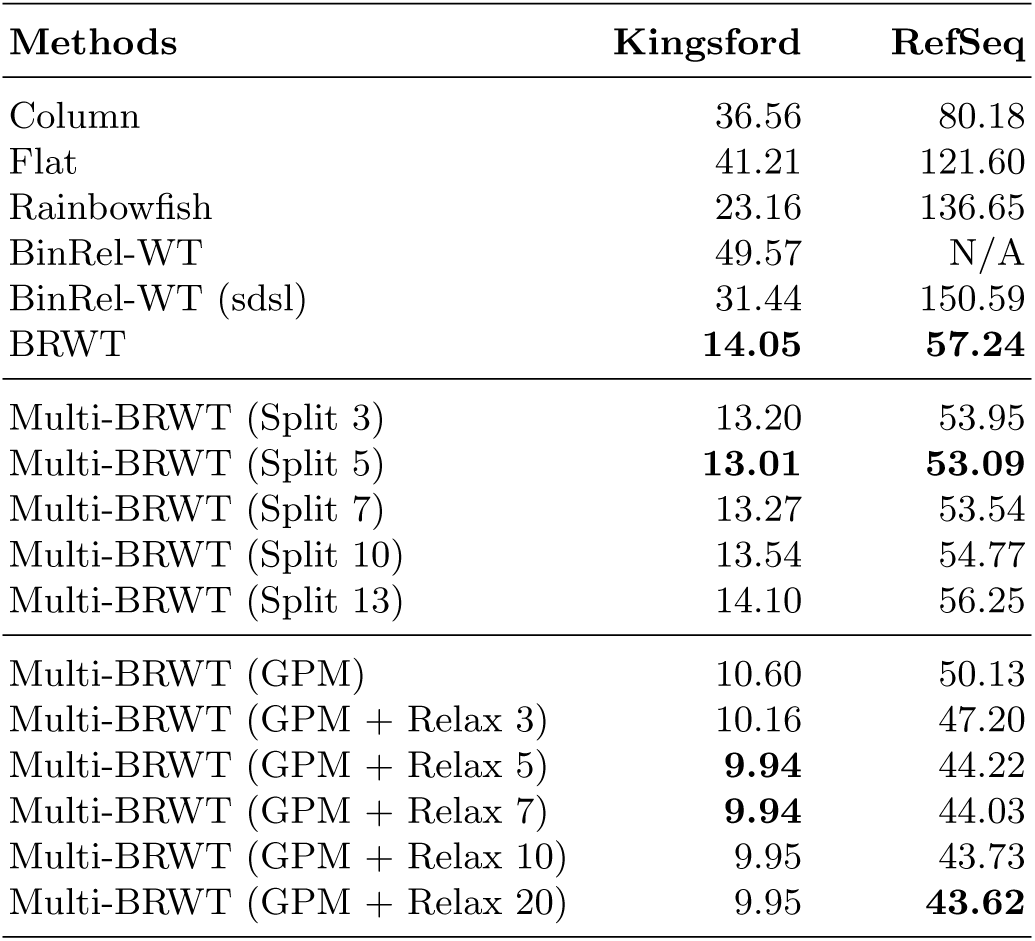
The measured size of the compressed binary relation matrix for different representations, in Gigabytes (Gb).

We improved the compression performance of a binary BRWT through the greedy pairwise matching (GPM) procedure described in Section 2.3. This strategy further decreases the size by another 18% to 10.6Gb. Finally, optimizing the tree topology using the GPM procedure and selectively removing internal nodes (reassigning children to their grandparents) while maintaining a constraint on each node’s maximum number of children, leads to the smallest space achieved in our experiments. By applying this technique, we decrease the required space to 9.94Gb (Multi-BRWT (GPM + Relax 5), with at most 5 children for each node). This is a 29% improvement over the basic BRWT representation and a 57% improvement over Rainbowfish. The Multi-BRWT (GPM + Relax 5) representation took 187 and 141 minutes of the compute time with 30 threads for the first and the second stages of the construction algorithm, respectively.

#### RefSeq Reference Genomes

Compression of the complete RefSeq genome annotation (release 88) resulted in a de Bruijn graph of dimension *k* = 15 containing *n* = 1, 073, 741, 824 nodes, leading to a binary relation matrix of *n* rows and *m* = 3, 173 columns with density ~ 3.8%, which is relatively high for a genome graph annotation and can be explained by the small *k*-mer size used.

This is a substantially larger dataset with less dependency between labels (columns). With the Multi-BRWT (GPM + Relax 20) representation, we were able to achieve a compressed storage size of only 43.6Gb (Table 1). Conversion from the column-compressed representation to Multi-BRWT (GPM + Relax 20) took 625 and 733 minutes of the compute time with 30 threads for the first and the second Multi-BRWT construction stages, respectively, which is quite reasonable for a real-world setting.

Also here, the basic BRWT method improves drastically over the column compressed baseline (29%), and the Multi-BRWT approach considerably surpasses the basic BRWT method (24% reduction in size). One can see that the state-of-the-art method Rainbowfish performs very poorly on the RefSeq dataset, which can be explained by the high density of the annotation matrix.

The construction of the BinRel-WT representation exceeded our available RAM (2Tb).

All experiments were performed on a Intel(R) Xeon(R) CPU E7-8867 v3 (2.50GHz) processor from ETH’s shared high-performance compute systems.

#### Supplementary Results

Compression ratios for methods using RRR vectors of block size 127 can be found in the Supplementary Materials.

## 4 Conclusion

We have presented a series of compressed representation methods for binary relations, building upon and improving on the existing literature. By generalizing BRWTs to multiary trees with improved partitioning schemes and adaptive arity to reduce data representation overhead, we have improved on state-of-the-art compression techniques for both simulated and real-world biological datasets.

We have shown that the structure of the input data has a strong influence on the compression performance and methods such as Rainbowfish benefit from presence of redundancy in rows or their correlations (when multiple objects carry a similar set of labels). It is noteworthy that in a real-world setting, where more and more labels are added to the set, the number of redundant rows decreases (ultimately leading to a set of mostly independent rows) and these methods work less well. Interestingly, it is especially this setting that regularly occurs in the labeling of genome graphs, where an underlying set of (related) sequences is assigned a growing set of different labels.

We have presented a method that copes very well with an increasing number of related columns as well as with the increasing density of the compressed binary matrix, and we showed that this results in considerable performance gains on both synthetic and typical real-world data. Our method, Multi-BRWT, led to a 24-29% reduction in size compared to the basic BRWT scheme on real-world data, and to a 57-68% reduction compared to the closest state-of-the-art method for compressing graph annotations, Rainbowfish.

A natural extension of this work will involve the utilization of dynamic vectors in the underlying storage of BRWTs to allow for their use in dynamic database contexts. Of particular interest are the ability to rearrange columns and use of dynamic compressed structures to avoid expensive decompression and recompression steps when performing updates.

Another interesting direction is the development of hybrid BRWT schemes that take the shape of Multi-BRWT but assign multiple columns to the leaves of the tree, using arbitrary schemes for compressing these. This would take advantage of both column and row structure in the binary relation matrix. These approaches are also beneficial for tackling the problem of achieving similar time complexities for both object and label queries on the compressed representation of the binary relations.

Overall, we conclude that, despite the advancements in compression over the recent years, there is still much room and many degrees of freedom in compressor design for further improvement.

## Supplementary Materials

Supplementary materials may be accessed via the bioRxiv pre-print located at https://doi.org/10.1101/468512.

## Acknowledgements

We would like to thank the members of the Biomedical Informatics group for fruitful discussions and critical questions, and Torsten Hoefler and Mario Stanke for constructive feedback on the graph setup. Harun Mustafa and Mikhail Karasikov are funded by the Swiss National Science Foundation grant #407540 167331 “Scalable Genome Graph Data Structures for Metagenomics and Genome Annotation” as part of Swiss National Research Programme (NRP) 75 “Big Data”.

