## Supplementary Material for "Sparse Binary Relation Representations for Genome Graph Annotation"

### 1 Binary relation matrix simulation

To benchmark our compression techniques systematically, we generated three series of random binary matrices satisfying different properties. Given fixed matrix dimensions $n \times m$, an expected column density $d$, and a uniqueness factor $u$, we define our generation schemes as follows:

1. **Random:** generate $m$ random columns of length $n$ with expected density $d$
2. **Uniform rows:** generate $m$ random columns of length $\frac{n}{u}$, then duplicate each row $u$ times
3. **Uniform columns:** generate $\frac{m}{u}$ columns of length $n$, then duplicate each column $u$ times

For each generated column, its indices are iterated through linearly and the values of the indices are set by drawing observations from a random variable $X \sim \text{Bernoulli}(d)$. For all experiments, values of $n = 1,000,000$, $u = 5$, and $d = 0.01$ were used. The values $m \in \{500, 1000, 3000\}$ were used.

#### 1.1 Sizes of compressed representations

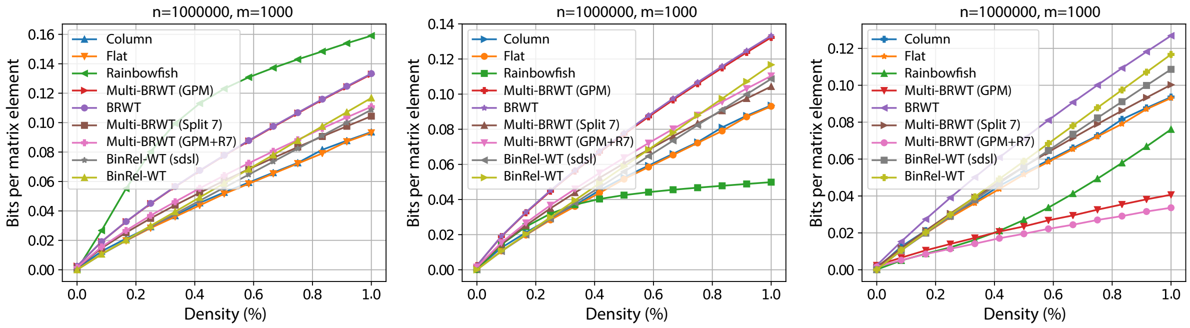


**Fig. 1.** Size of the representation of $A \in \{0,1\}^{10^6 \times 1000}$ with densities $d < 0.01$ using different approaches: a) Random columns; b) Duplicated rows; c) Duplicated columns.

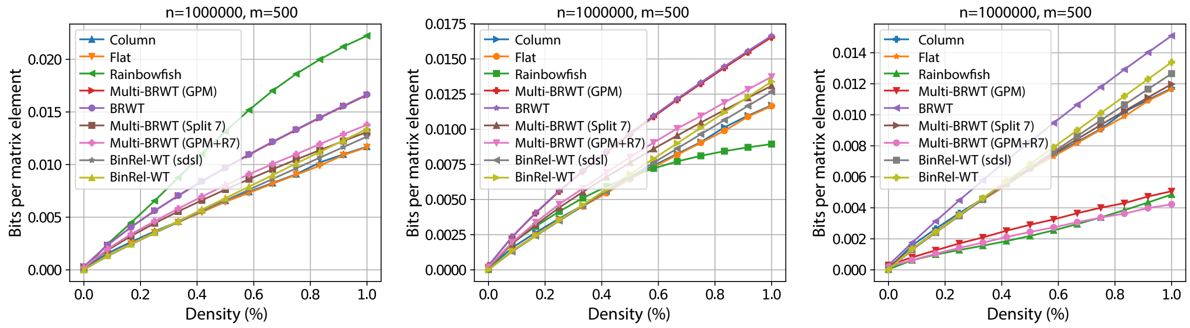


**Fig. 2.** Size of the representation of $A \in \{0,1\}^{10^6 \times 500}$ with densities $d < 0.01$ using different approaches: a) Random columns; b) Duplicated rows; c) Duplicated columns.

### 2 Subsampling lemma

*Proof (Subsampling lemma).* For each $i \in \{1, ..., n\}$, we introduce a random variable $X_i$ as follows:

$$X_i = \begin{cases} 1 & \text{if } i \in S, \\ 0 & \text{otherwise.} \end{cases}$$

In other words, $X_i$ is 1 if $i$ is subsampled (i.e., $S \ni i$), and 0 otherwise. In addition, for each subset $V \subset \{1, ..., n\}$, we define $X_V = \sum_{i \in V} X_i = |V \cap S|$, which counts how many elements of $V$ are subsampled. Thus,

$$X_{o_i \cup o_j} = |(o_i \cup o_j) \cap S| = |\tilde{o}_i \cup \tilde{o}_j| = p\, \hat{u}_{ij},$$
$$\mathbb{E}X_{o_i \cup o_j} = |o_i \cup o_j| \cdot \mathbb{E}X_1 = |o_i \cup o_j| \cdot p = p\, u_{ij}.$$

Now we derive

$$\begin{aligned} \Pr(|\hat{u}_{ij} - u_{ij}| \geq \varepsilon u_{ij}) &= \Pr\Big( \Big| \frac{1}{p} X_{o_i \cup o_j} - \frac{1}{p} \mathbb{E}X_{o_i \cup o_j} \Big| \geq \frac{1}{p} \varepsilon \mathbb{E}X_{o_i \cup o_j} \Big) \\ &= \Pr\Big( \big| X_{o_i \cup o_j} - \mathbb{E}X_{o_i \cup o_j} \big| \geq \varepsilon \mathbb{E}X_{o_i \cup o_j} \Big), \end{aligned}$$

and use the Chernoff bound for $\varepsilon \in (0, 1)$

$$\Pr\Big( |X_{o_i \cup o_j} - \mathbb{E}X_{o_i \cup o_j}| \geq \varepsilon \mathbb{E}X_{o_i \cup o_j} \Big) \leq 2e^{-\varepsilon^2 \mathbb{E}X_{o_i \cup o_j}/3} = 2e^{-\varepsilon^2 p u_{ij}/3} \leq 2e^{-\varepsilon^2 pd/3}. \qquad (1)$$

Therefore, we can bound the joint probability as follows:

$$\begin{aligned} \Pr\Big( \bigcap_{i,j=1}^{m} \Big\{ |\hat{u}_{ij} - u_{ij}| < \varepsilon u_{ij} \Big\} \Big) &= 1 - \Pr\Big( \bigcup_{i,j=1}^{m} \Big\{ |\hat{u}_{ij} - u_{ij}| \geq \varepsilon u_{ij} \Big\} \Big) \\ &\geq 1 - \sum_{i \leq j} \Pr(|\hat{u}_{ij} - u_{ij}| \geq \varepsilon u_{ij}) && \triangleright \text{ union bound} \\ &\geq 1 - \frac{m^2 + m}{2} \cdot 2e^{-\varepsilon^2 pd/3}. && \triangleright \text{ by inequality (1)} \end{aligned}$$

Finally, we reformulate this result in the following equivalent form:

$$\Pr\Big( \bigcap_{i,j=1}^{m} \Big\{ |\hat{u}_{ij} - u_{ij}| < \varepsilon u_{ij} \Big\} \Big) \geq 1 - \delta,$$

if $\delta \geq (m^2 + m) e^{-\varepsilon^2 pd/3}$ or, equivalently, if

$$p \geq \frac{3 \ln(\frac{m^2+m}{\delta})}{d \varepsilon^2}.$$

**Table 1.** The measured size of the compressed binary relation matrix for different methods in Gigabytes. A block size of 127 was used for the underlying RRR vectors.

| **Methods** | **Kingsford** | **RefSeq** |
|---|---|---|
| Column | 36.56 | 80.18 |
| Flat | 41.21 | 121.60 |
| Rainbowfish | 19.22 | 117.31 |
| BinRel-WT | 49.57 | N/A |
| BinRel-WT (sdsl) | 31.40 | 147.03 |
| BRWT | **12.97** | **51.82** |
| Multi-BRWT (Split 3) | 12.24 | 49.29 |
| Multi-BRWT (Split 5) | **12.01** | 48.25 |
| Multi-BRWT (Split 7) | 12.13 | **48.18** |
| Multi-BRWT (Split 10) | 12.28 | 48.65 |
| Multi-BRWT (Split 13) | 12.61 | 49.36 |
| Multi-BRWT (GPM) | 9.68 | 48.10 |
| Multi-BRWT (GPM + Relax 3) | 9.36 | 45.45 |
| Multi-BRWT (GPM + Relax 5) | **9.19** | 42.75 |
| Multi-BRWT (GPM + Relax 7) | 9.21 | 42.56 |
| Multi-BRWT (GPM + Relax 10) | 9.22 | 42.34 |
| Multi-BRWT (GPM + Relax 20) | 9.22 | **42.28** |
